# Generation of Transgenic Human Malaria Parasites With Strong Fluorescence in the Transmission Stages

**DOI:** 10.1101/500793

**Authors:** Kyle McLean, Judith Straimer, Christine S. Hopp, Joel Vega-Rodriguez, Abhai Tripathi, Godfree Mlambo, Peter C. Doumolin, Chantal T. Harris, Xinran Tong, Melanie J. Shears, Johan Ankarklev, Björn F.C. Kafsak, David A. Fidock, Photini Sinnis

**Affiliations:** Department of Molecular Microbiology and Immunology, and Johns Hopkins Malaria Institute, Johns Hopkins Bloomberg School of Public Health, Baltimore, MD 21205; Department of Microbiology and Immunology, Columbia University Irving Medical Center, New York, NY 10032; Division of Infectious Diseases, Department of Medicine, Columbia University Irving Medical Center, New York, NY 10032; Department of Microbiology and Immunology, Weill Cornell School of Medicine, New York, NY 10065

## Abstract

Malaria parasites have a complex life cycle that includes specialized stages for transmission between their mosquito and human hosts. These stages are an understudied part of the lifecycle yet targeting them is an essential component of the effort to shrink the malaria map. The human parasite *Plasmodium falciparum* is responsible for the majority of deaths due to malaria. Our goal was to generate transgenic *P. falciparum* lines that could complete the lifecycle and produce fluorescent transmission stages for more in-depth and high-throughput studies. Using zinc-finger nuclease technology to engineer a marker-free integration site, we generated three transgenic *P. falciparum* lines in which *tdtomato* or *gfp* were stably integrated into the genome. Expression was driven by either stage-specific *peg4* and *csp* promoters or the constitutive *ef1a* promoter. Phenotypic characterization of these lines demonstrates that they complete the life cycle with high infection rates and give rise to fluorescent mosquito stages. The transmission stages are sufficiently bright for intra-vital imaging, flow cytometry and scalable screening of chemical inhibitors and potentially inhibitory antibodies.

## Introduction

Malaria remains one of the most important infectious diseases in the world, impacting approximately a third of the world’s population and responsible for over 400,000 deaths annually [1]. The disease is caused by parasites of the genus *Plasmodium*, transmitted to their mammalian host by the infected Anopheline mosquitoes. Malaria parasites have two distinct stages in their mammalian hosts: an asymptomatic pre-erythrocytic stage, consisting of sporozoites and the liver stages into which they develop, and an erythrocytic stage when high parasite numbers lead to clinical disease. A small proportion of blood stage parasites differentiate to gametocytes which, when taken up in the blood meal, initiate infection in the mosquito, giving rise to oocysts on the midgut wall that produce sporozoites that migrate to the mosquito’s salivary glands. The transmission stages of *Plasmodium*, namely sporozoites and gametocytes, are bottlenecks in the parasite life cycle and their low numbers has limited investigations into their biology and chemotherapeutic interventions.

Rodent malaria models have enabled more in-depth study of the difficult to access transmission stages and have played a particularly important role in our understanding of pre-erythrocytic stages. Nonetheless, there are species-specific differences across the *Plasmodium* genus, such as the unique morphology and prolonged development of gametocytes of the *Laverania*, that make it critical to perform confirmatory studies and inhibitor screening with human malaria parasites. In order to facilitate these investigations, we generated transgenic *Plasmodium falciparum* lines expressing tdTomato or Green Fluorescent Protein (GFP) under the control of pre-erythrocytic or gametocyte stage promoters. Here we describe the generation of these parasites and their fluorescent properties throughout the development of the transmission stages.

## Results and Discussion

### Generation of fluorescent *P. falciparum* lines

Previous studies have generated fluorescent *P. falciparum* lines using the constitutively expressed elongation factor *eEF1α* (PF3D7_1357000) promoter and Green Fluorescent Protein (GFP) variants [2] [3]. While useful, the low intensity of fluorescence of these parasites during sporozoite and exoerythrocytic stages has limited their utility, particularly when imaging in mosquito and mammalian tissues where there is a high degree of autofluorescence in the green channel. We reasoned that using the fluorescent protein tandem-dimer Tomato (tdTomato) [4] in the place of GFP could surmount some of these challenges, as it is much brighter and matures faster than GFP [5] and the red-shifted fluorescence is better suited for intravital tissue imaging.

Additionally, we reasoned that strong, stage-specific promoters could be used to further boost the fluorescent signal during the sporozoite and exoerythrocytic stage. We tested two potential candidates, the promoter of the circumsporozoite protein (CSP) gene (PF3D7_0304600), which drives expression of the most abundant sporozoite surface protein, and the promoter of the Plasmodium Early Gametocyte 4 (PEG4, also known as eTRAMP10.3) (PF3D7_1016900) gene. PEG4 was first described as a protein expressed in the early stages of gametocytogenesis [6], and its promoter region has been shown to be active at those times [2]. Indeed, the utility of this promoter to drive episomal transgene expression in *P. falciparum* gametocytes has been previously demonstrated [2]. However, fluorescence in mosquito stages was never assessed as these studies used a line that does not make infection-competent gametocytes. More recently, PEG4 was reported to be the homolog of the well-studied *P. yoelii upregulated in sporozoites 4* (UIS4) gene [7]. The *uis4* promoter has been a valuable tool for driving strong fluorescence in sporozoites of *P. yoellii and P. berghei* [8] and we hypothesized that the same would be true for *P. falciparum*, potentially providing a promoter that could drive expression in both sporozoites and gametocytes.

Typically, generation of genetically-modified *P. falciparum* parasites requires long *in vitro* selection and cloning times, which frequently results in the loss of the parasite’s ability to generate gametocytes or infect the mosquito. Beginning with a highly mosquito-infectious clone of the *P.falciparum* NF54 isolate [9], we used zinc-finger nuclease (ZFN) technology to rapidly engineer a selectable marker-free *Bxb1* phage integration site *(attB)* into the parasite genome (Figure 1A). We selected the coding sequence of the *pfs47* gene for *attB* introduction. Though recent work showed that Pfs47 is required for parasite evasion of the mosquito complement system in *Anopheles gambiae* mosquitoes [10], it is not required for parasite survival in *Anopheles stephensi* [11] [12] and had been used as a locus for transgene expression in mosquito-stages by others [3] [13]. Once a stably-transfected parasite population was obtained, the line was cloned by limiting dilution, and the *pfs47* locus was sequenced to verify the presence of the *attB* site (Figure 1B). Clones were subsequently tested for gametocyte production and mosquito infectivity.

**Figure 1.**
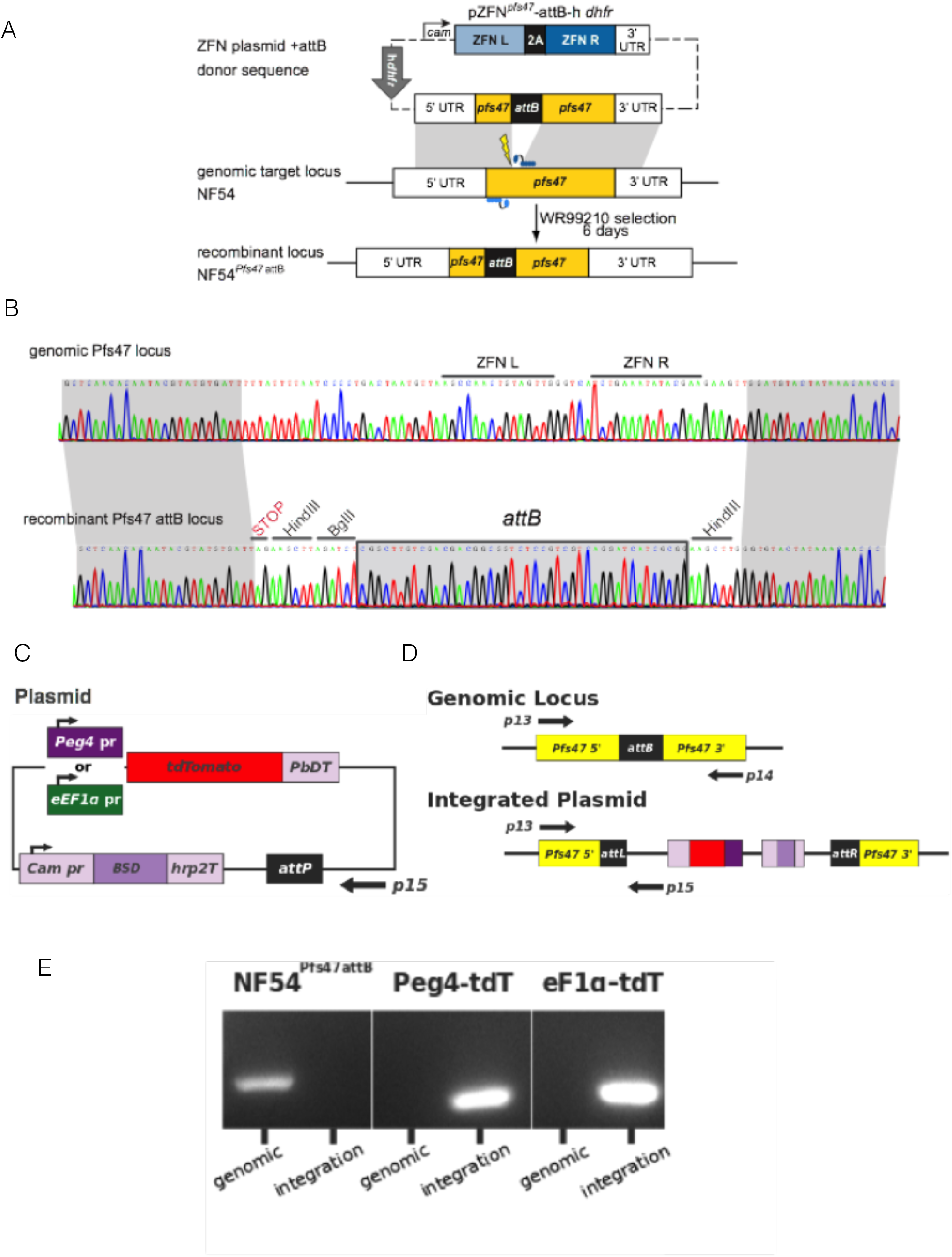
Introduction of the *attB* site into *pfs47* locus and integration of tdTomato transgenes. A) Schematic of the *attB* integration strategy. The expression of the *pfs47*-specific ZFN pair is under the control of the *calmodulin* promoter and the *hsp86* 3’UTR. The homologous donor sequence provided for double-strand break repair comprises a fragment of *pfs47* stretching 0.68 kb upstream and 0.87 kb downstream of the ZFN target site (thunderbolt). The attB site is located between the two homologous regions and will replace the genomic *pfs47* sequence containing the ZFNs binding site upon integration. The transfection plasmid pZFN*^pfs47^*-attB-*hdhfr* also contains a selectable marker for transient selection and propagation of the plasmid. B) Sequence analysis of the unmodified endogenous *pfs47* locus of the parental parasite line NF54 and the gene-edited recombinant locus NF54^pf47-attB^. C) Schematic of the *tdTomato* plasmids used for transfection *(PbDT = P. berghei* DHFR-TS terminator, *Cam pr = calmodulin* promoter, *BSD* = Blasticidin S Deaminase selectable marker, *hrp2T* = HRP2 terminator). D) Schematic of the NF54^Pfs47attB^ *attB* locus pre- and post-integration of the *tdTomato* plasmids. Position of primers used to amplify the genomic locus (p13 & p14) and the integrated plasmid (p13 & p15) are shown. E) PCR verification of plasmid integration. Primers p13 and p14 generate a 334 bp product from the unmodified *pfs47attB locus* (genomic), and primers p13 and p15 generate at 287 bp product upon *tdTomato* plasmid integration (integration).

A clone with high production of infectious gametocytes (NF54^Pfs47attB^C10) was then selected for secondary transfection. Using the *Bxb1* integrase system [14], a plasmid containing either Pf-*ef1α*-tdTomato, Pf-*peg4*-tdTomato, (Figure 1C) or Pf-*csp*-GFP expression cassettes along with a Blasticidin S Deaminase (BSD) resistance cassette was introduced into the chromosomal *pfs47 attB* site. Stably transfected cultures were cloned by limiting dilution and tested for plasmid integration (Figure 1D) and gametocyte production. Three clones of each of the tdTomato lines were passed through the mosquito to identify a clone producing robust sporozoite numbers. Pf-*peg4*-tdTomato Clone A and Pf-*ef1α*-tdTomato Clone 3D06 were chosen for further characterization. The Pf-*csp*-GFP line was not cloned and the parental line was studied.

### Phenotypic characterization of *P. falciparum* fluorescent lines

To characterize the tdTomato clones and the uncloned GFP line in the mosquito host, *Anopheles stephensi* mosquitoes were fed with human blood containing gametocytes and subsequent parasite development in mosquitoes was followed. Oocyst numbers for the tdTomato lines were robust and as expected, both Pf-*ef1α*-tdTomato and Pf-*peg4*-tdTomato oocysts were fluorescent (Figure 2A&B). Indeed at this stage, Pf-*ef1α*-tdTomato fluorescence is quite strong, likely because each oocyst consists of many genomes. Oocyst numbers in the GFP line were also within the range normally observed in the parental NF54 line (Figure S1).

**Figure 2.**
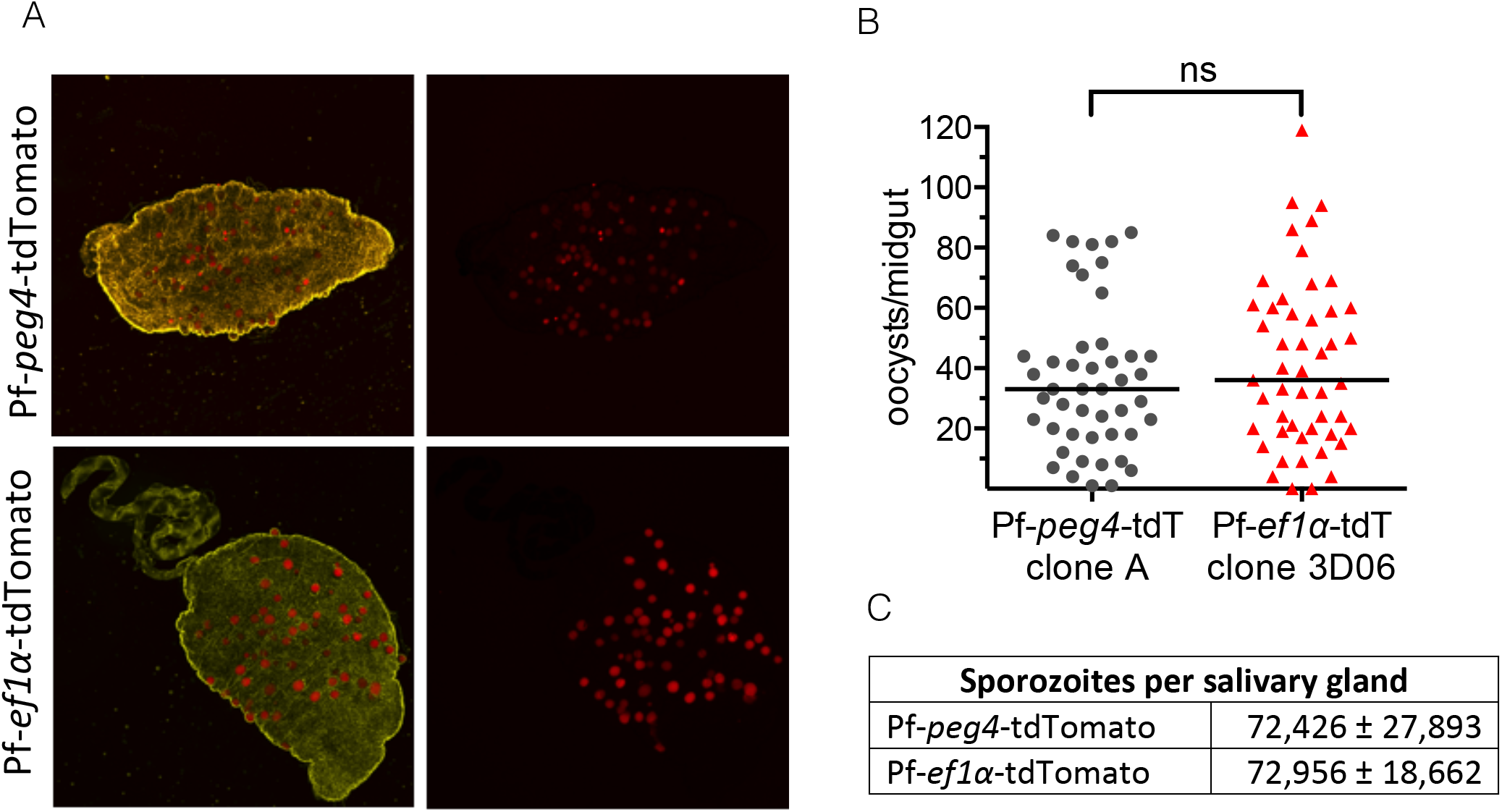
Passage through the mosquito of *P. falciparum*-tdTomato lines is normal. A) Mosquito midguts (whitelight) were dissected 10 days after the infectious blood meal and imaged using fluoescence microscopy to visualize associated Pf-*peg4*-tdTomato and Pf-*ef1 α-*tdTomato oocysts (red). B) Oocysts were counted based on their tdTomato fluorescence and data from two independent feeding experiments each are shown. Horizontal line shows median oocyst number. C) Pf-*peg4*-tdTomato and Pf-*ef1α*-tdTomato lines produce normal number of salivary gland sporozoites. Shown is the mean +/- standard deviation of salivary gland loads from 20 mosquitoes.

In all three lines, oocyst sporozoites progressed to invade salivary glands with sporozoite numbers similar to those routinely observed in the NF54 parental line (Figures 2C and 3C). As expected, salivary glands infected with Pf-*ef1α*-tdTomato sporozoites were dimly fluorescent, while Pf-*peg4*-tdTomato infected salivary glands were brightly fluorescent and could easily be visualized through the chitin exoskeleton of the mosquito (Figure 3A), confirming that the *peg4* promoter is a strong sporozoite promoter. We then performed immunofluorescence assays on salivary gland sporozoites to compare the endogenous tdTomato or GFP fluorescence with the signal observed upon immunostaining of the major surface protein, the circumsporozoite protein (CSP). The tdTomato signal when under the control of the *peg4* promoter was similar to the strong CSP signal while sporozoites expressing tdTomato under the *ef1α* promoter were only faintly fluorescent (Figure 3B). The GFP-transgenic sporozoites were similarly strongly fluorescent, as might be expected given the strength of the *csp* promoter at this stage (Figure S1, panel D).

**Figure 3.**
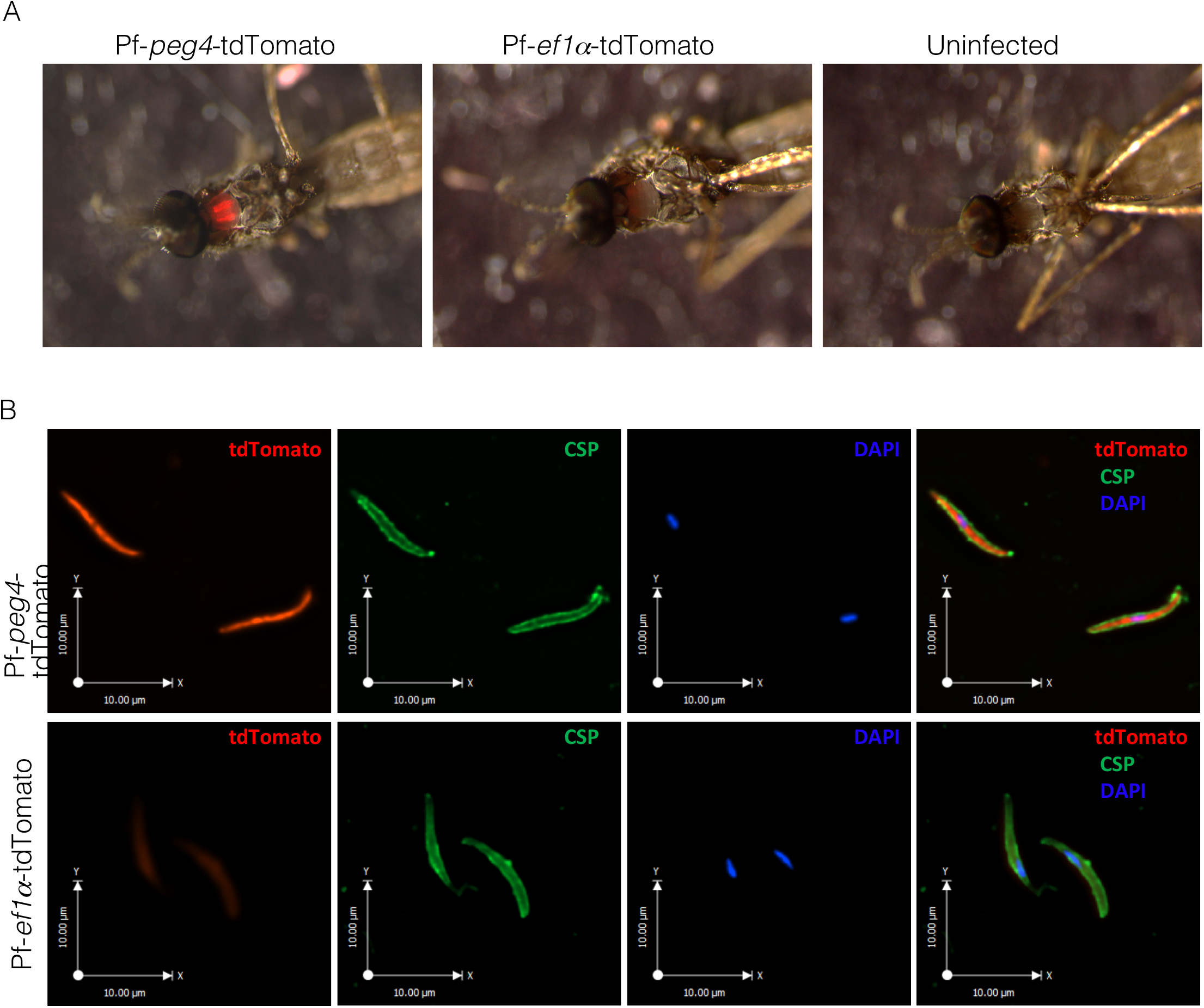
Immunofluorescence of sporozoites of *P. falciparum-tdTomato* lines. A) The thorax of mosquitoes infected with the Pf-*ef1α*-tdTomato or Pf-*peg4*-tdTomato lines was visualized under a fluorescence stereoscope to detect the presence of the tdTomato sporozoites in the salivary glands. Note the red fluorescence observed in the thorax of mosquitoes infected with the *P. falciparum-tdTomato* lines compared to the absence of fluoresce in the uninfected mosquito. B) Salivary gland sporozoites from mosquitoes infected with the Pf-*ef1α*-tdTomato or Pf-*peg4*-tdTomato were fixed and stained with an anti-PfCSP antibody (green) and analyzed by fluorescence microscopy to detect the cytoplasmic tdTomato (red) and the surface CSP (green). DNA was stained with DAPI (blue). Note the more intense signal detected from Pf-*peg4*-tdTomato sporozoites as compared to Pf-*ef1α*-tdTomato sporozoites, which match the fluorescence intensities from panel A.

We also investigated the fluorescence of the transgenic tdTomato lines in the liver stage of the parasite by performing *in vitro* infection assays using sporozoites and primary human hepatocytes. Immunofluorescence assays of liver stage parasites co-stained with an mAb to heat shock protein 70 (hsp 70) and DAPI to visualize the DNA, showed that both the Pf-*ef1α*-tdTomato and Pf-*peg4*-tdTomato lines were brightly fluorescent in this stage (Figure 4A). Furthermore, live imaging confirmed that liver stage parasites were brightly fluorescent and could be clearly seen without immunostaining (Figure 4B). These data indicate that both lines could be used to study and isolate *P. falciparum* liver stage parasites.

**Figure 4.**
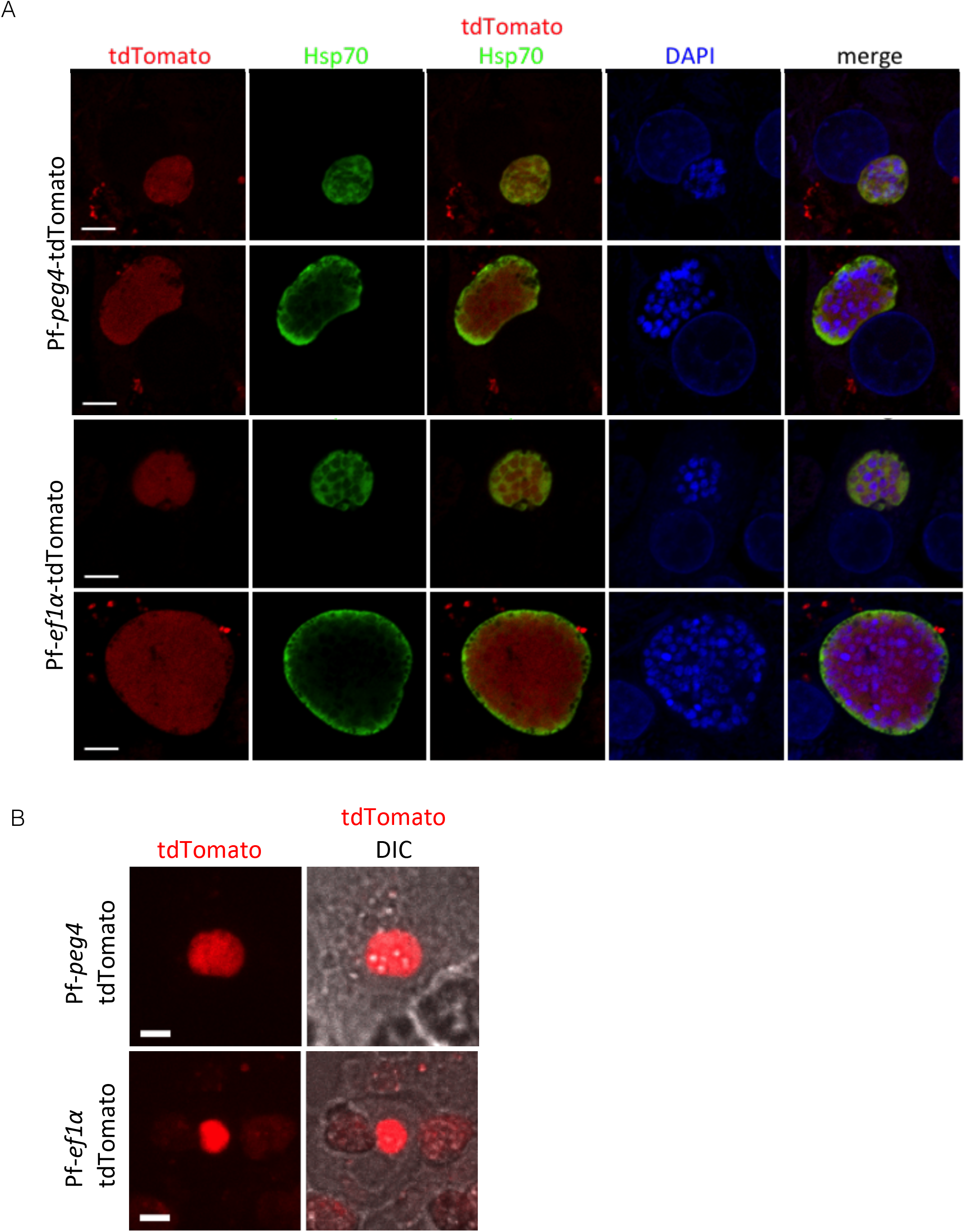
Immunofluorescene assays of late liver stage *P. falciparum-tdTomato* lines. A) Primary human hepatocytes were infected with Pf-*peg4*-tdTomato and Pf-*ef1α*-tdTomato sporozoites. At 96 hours post infection, cells were fixed and stained for PfHsp70 (green) to visualize the parasite cytoplasm together with the endogenous tdTomato fluorescence (red). DNA was stained with DAPI (blue). Scale bars 5 μm. B) Live confocal microscopy of Pf-*peg4*-tdTomato and Pf-*ef1α*-tdTomato EEFs at 96 hours post infection of primary human hepatocytes. Fluorescent images and overlays with the DIC images are shown. Scale bars 15 μm.

As mentioned above, PEG4 is likely the *P. falciparum* ortholog of the rodent malaria parasite protein UIS4 [7], whose expression is highly upregulated in sporozoites and early liver stages [15] [16]. In contrast to the rodent parasites, *P. falciparum* PEG4 is also expressed in gametocytes [6]. We therefore quantified the fluorescence of Pf-*peg4*-tdTomato gametocytes over the entire course of their development. Using flow cytometry to obtain fluorescence data from large numbers of gametocytes we observed a strong red fluorescence signal above asexual parasite background beginning with Stage I gametocytes (day 2 of maturation). The fluorescence increased substantially from stage III gametocytes (day 5) onward and reached a peak of expression in mature stage V gametocytes (day 12) (Figure 5A). Images of gametocytes over the course of their development illustrate the expression of the tdTomato transgene in the different gametocyte stages of the Pf-*peg4*-tdTomato line (Figure 5B). Furthermore, exflagellation of male gametocytes could also be visualized (Figure 5C). Thus this parasite line will be useful to laboratories studying the sexual stages of *P. falciparum* and could streamline drug screening targeting gametocyte development.

**Figure 5.**
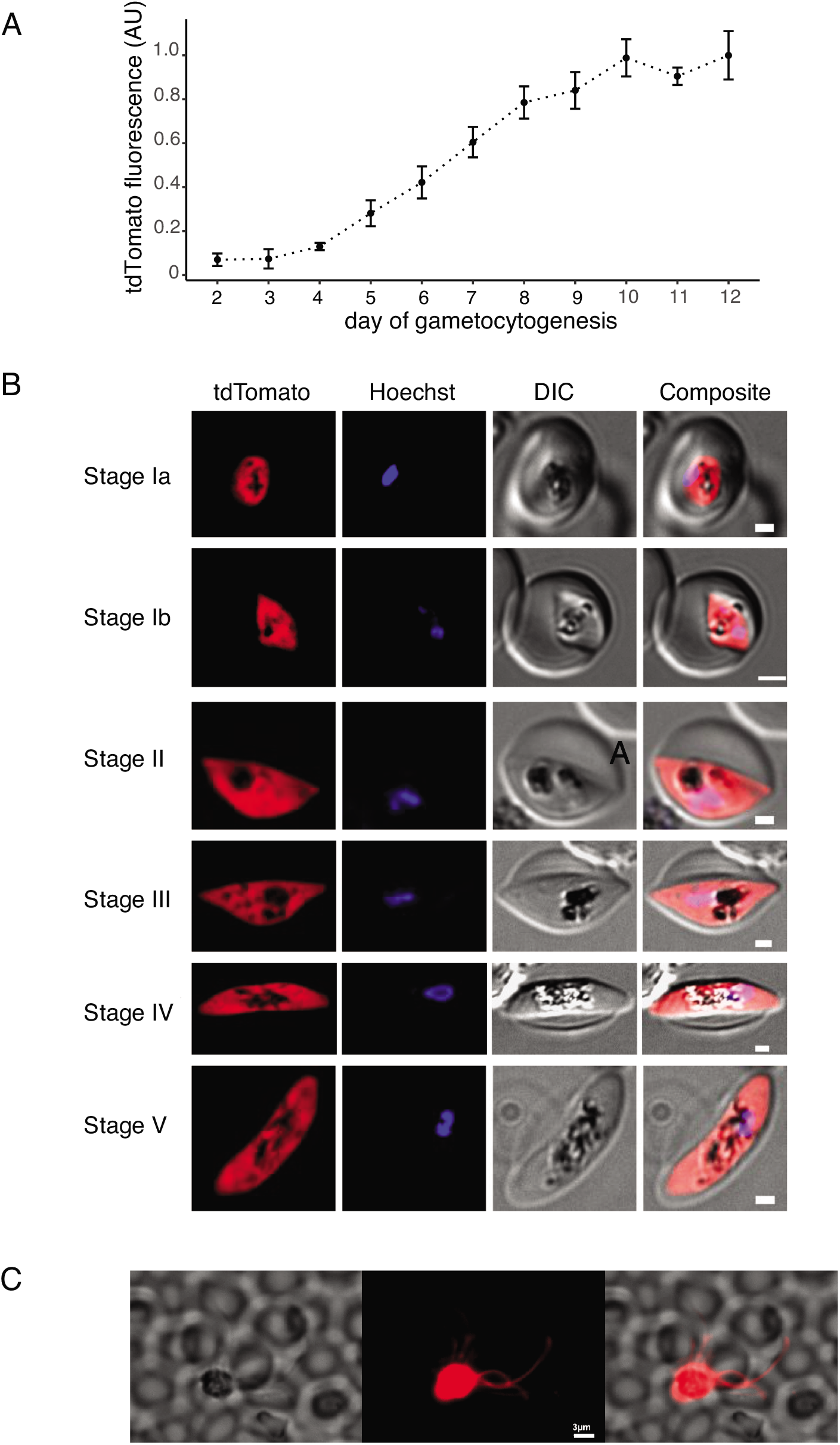
tdTomato expression increases throughout Pf-*peg4*-tdTomato gametocyte development. A) Mean fluorescence of >1000 gametocytes was measured on days 2-12 by flow cytometry. Values are normalized to the maximum mean tdTomato signal on day 12. mean +/- standard deviation from two independent experiments is shown. B) Representative phase and fluorescence images of Pf-PEG4-tdTomato stage I-V gametocytes with the nuclear stain Hoechst 33342. Scale bar = 1 micron. C) Exflagellating Pf-*peg4*-tdTomato microgametes. Scale bar = 3 microns.

In conclusion, we have generated three transgenic fluorescent *P. falciparum* lines that robustly complete the parasite life cycle. The Pf-*peg4*-tdTomato and Pf-*csp*-GFP lines produce highly fluorescent sporozoites that are sufficiently bright to be used for intravital imaging and could be used in combination with humanized mice [17] to gain information on the behavior of human malaria sporozoites *in vivo*. To date, these types of studies have only been performed with rodent malaria parasites [8] [18]. Furthermore, our data demonstrate that the *peg4* promoter is a strong sporozoite-stage promoter in the human malaria parasite *P. falciparum*. The Pf-*peg4*-tdTomato line additionally produces highly fluorescent gametocytes, which can be quantified by flow cytometry to more rapidly screen inhibitors of gametocytogenesis and thus identify compounds that can inhibit transmission of the parasite to the mosquito. Though this has been previously demonstrated using a *P. falciparum* line episomally expressing tdTomato under the control of the *peg4* promoter [2], there are advantages to using lines with stably-integrated transgenes as the expression levels from transgenes are more homogenous than is observed with episomal constructs that vary in copy number between parasites [14]. Furthermore, the gametocytes produced by our Pf-*peg4*-tdTomato line are infection-competent thus making transmission studies, the gold-standard of gametocyte maturation, feasible. Overall, these lines will enable more rapid screening of both chemical inhibitors and potentially inhibitory antibodies on transmission stages of *P. falciparum* and aid in flow sorting transmission stages for downstream analyses such as transcriptomic and proteomic studies.

## Methods

### Asexual Parasite Culture and Maintenance

Asexual blood stage parasites were propagated in 2% human erythrocytes in RPMI-1640 malaria culture medium containing 2 mM L-glutamine, 50 mg/L hypoxanthine, 25 mM HEPES, 0.225% NaHCO_3_, 10 mg/L gentamycin and 0.5% (w/v) Albumax II (Invitrogen). Parasites were maintained at 37°C under an atmosphere of 5% O_2_, 5% CO_2_ and 90% N_2_.

### Generation of the NF54^Pfs47attB^ *Plasmodium falciparum* line

Zinc-finger nucleases (ZFN) directed against the *P. falciparum 6-cysteine protein P47* (PlasmoDB ID PF3D7_1346800, also known as *Pfs47*) were purchased from Sigma Life Sciences. To introduce the *attB* site (from *Mycobacterium smegmatis*) into the *pfs47* locus in *P. falciparum*, we generated a transfection plasmid with *pfs47* specific ZFNs, a selectable human *dhfr* marker and a donor fragment consisting of two *pfs47* homologous sequences flanking the *attB* site. The left and right ZFNs were 2A-linked and placed under the regulatory control of the 5’ *calmodulin* (PF3D7_1434200) promoter and the 3’ *hsp86* (PF3D7_0708500) terminator [19]. The two homologous regions flanking the ZFN cut site were first amplified separately using the primer pairs p1 and p2 (0.68kb) and p3+p4 (0.87kb). Primer p2 and p3 also contained the *attB* site and the two fragments were then linked by “splicing by overlap extension” PCR. The donor sequence was added to the ZFN-containing plasmid using the restriction enzymes AatII and BstAPI, thus yielding the final pZFN^pfs47^-attB-h*dhfr* transfection plasmid. Asexual NF54 blood-stage parasites were electroporated with purified circular plasmid pZFN*^pfs47^*-attB-h*dhfr* as described (PMID: 9855645). One day after electroporation, parasites were exposed to 2.5 nM WR99210 for 6 days to select for transformed parasites. Integration of the *attB* site was assessed by PCR analysis of bulk cultures, and cultures showing evidence of editing were selected for 96-well cloning by limiting dilution. Clones were identified by staining with 2× SYBR Green I and 165 nM MitoTracker Deep Red (Invitrogen) and assaying for growth by flow cytometry on an Accuri C6 cytometer (Ekland *et al*., FASEB J. 2011 PMID: 21746861). To test for *attB* integration, *P. falciparum* genomic DNA was extracted and purified using DNeasy Blood kits (Qiagen). *Pfs47* sequences were examined by PCR-amplifying the genomic locus with primers p5+p6 flanking the *pfs47* plasmid donor sequence. These products were amplified from bulk cultures or parasite clones. Sequencing was performed with the internal p7 and p8 primers. Four clones were expanded to test their ability to infect mosquitoes and of these, NF54^Pfs47attB^ C10 gave the most robust mosquito infections and was therefore used for production of the fluorescent lines.

### Generation of fluorescent *P. falciparum* lines

Plasmid construction was performed using standard molecular cloning techniques. The *Pf-ef1α* used was the same 806bp sequence described and used by Talman *et al*. (PMID: 20161781), and was amplified by genomic DNA using primers p9 and p10. The 2227bp *Pf-peg4* upstream sequenced used was identical to that described by Buchholz *et al*. (PMID: 21502082), and was amplified from genomic DNA using primers p11 and p12. The *Pf-csp* sequence used consisted of a 1266bp sequence from upstream of the *csp* coding sequence (PF3D7_0304600). The final plasmids were denoted pBattP-eEF1apr_tdTomato, pBattP_PEG4pr_tdTomato, and pLN_CSpr-GFP (see *Supplement* for sequence files). Transfections were performed with 50μg of the fluorescent protein plasmid along with 50μg of the pINT integrase plasmid (genbank plasmid DQ813654.3) (PMID: 16862136) using the modified method described by Spalding *et al*. (PMID: 20403390). Transfections were selected with 2.5μg/mL Blasticidin S until stable parasites were obtained. Transfected lines were cloned by limiting dilution as described above. Chromosomal integration of the plasmids was verified by PCR using primers p13 and p15, and compared to PCR of the unmodified pfs47attB locus with primers p13 and p14.

### Gametocyte cultures and mosquito feeding

Gametocytes cultures were maintained as described above except with the following modifications: Culture medium contained 10% *v/v* human serum in the place of Albumax II, and parasites were propagated at 4% hematocrit. Instead of a gas incubator, cultures were maintained at 37°C in a candle jar made of glass desiccators. Gametocyte cultures were initiated at 0.5% asynchronous asexual parasitemia from low passage stock and maintained up to day 18 with daily media changes but without any addition of fresh erythrocytes. Day 15 to 18 cultures, containing largely mature gametocytes, were used for mosquito feeds: Cultures were centrifuged at 108 x *g* for 4 min and the parasite pellet was resuspended to final gametocytemia of 0.3% in a mixture of human O+ RBCs supplemented with 50% *v/v* human serum. Gametocytes were fed to *Anopheles stephensi* mosquitoes that had been starved overnight, using glass membrane feeders. Unfed mosquitoes were removed after feeding. *Anopheles stephensi* mosquitoes were then maintained on a 10% sugar solution at 25°C and 80% humidity with a 14:10 hr light:dark cycle including a 2 hr dawn/dusk transition. according to standard rearing methods. Human erythrocytes used to set up the cultures are collected weekly from healthy donors under an institutional review board-approved protocol.

### Oocyst and salivary gland sporozoite quantification

*An. stephensi* mosquitoes infected with Pf-*ef1α*-tdTomato or Pf-*peg4*-tdTomato were maintained at 25°C, 80% humidity and on days 10 and 14 after the infective-blood meal, mosquitoes were dissected and midguts or salivary glands were harvested for sporozoite counts. On day 10 midguts were observed and photographed for oocyst counts by fluorescence and phase microscopy using an upright Nikon E600 microscope with a PlanApo 10× objective. On day 14, salivary glands from 30-50 mosquitoes were pooled, and released sporozoites were counted using a haemocytometer. For direct visualization of Pf-*ef1α*-tdTomato or Pf-*peg4*-tdTomato sporozoites in situ, infected mosquitoes were immobilized at 4°C and transferred to a petri dish pre-chilled on ice. Mosquitoes were observed under an Olympus SZX7 fluorescence stereo microscope for detection of red fluorescence from transgenic sporozoites that had invaded the salivary glands inside the mosquito thoracic cavity. Images of the mosquitoes were obtained using the Q-Capture Pro 7 software. Uninfected mosquitoes were used as negative control.

### Immunofluorescence assays (IFAs)

Sporozoites. Pf-*ef1α*-tdTomato or Pf-*peg4*-tdTomato sporozoites were isolated from infected mosquitoes 16 days post-infection. Sporozoites were centrifuged onto a glass coverslip at 300 x g for 4 min to increase attachment to the coverslip. Sporozoites were fixed with 4% paraformaldehyde for 10 min, washed with PBS and then incubated with Alexa Fluor 488 labeled mAb 2A10, specific for the *P. falciparum* circumsporozoite protein (CSP; REF) diluted in PBS to a concentration of 1 μg/ml for 1 hour at room temperature. After three washes with PBS, sporozoites were mounted with ProLong Gold Antifade with DAPI (Thermofisher) and visualized on a Zeiss Axio Imager M2 fluorescence microscope for detection of tdTomato red fluorescence, PfCSP green fluorescence and DAPI. Sporozoite images were obtained and analyzed using the Volocity software.

Exoerythrocytic Stages. Cryopreserved human primary hepatocytes, hepatocyte thawing, plating and maintenance medium were obtained from Triangle Research Labs (TRL, North Carolina). Growth and infection of primary hepatocytes was performed as previously outlined in (Dumoulin et al. 2015 Plos One). Briefly, primary hepatocytes (donor 4051) were thawed and 300,000 hepatocytes were plated on collagen-coated coverslips in a 24 well plate. 48 hrs later, 3-5 x 10^5^ Pf-*ef1α*-tdTomato or Pf-*peg4*-tdTomato sporozoites were added per well in Iscove’s Modified Dulbecco’s Medium (IMDM) with 2.5% FCS supplemented with penicillin, streptomycin and L-glutamine (complete growth medium), centrifuged at 380 x *g* for 5 min without break and then incubated at 37°C for 4-6 hours. After incubation, cells were washed with PBS and maintained in complete growth medium supplemented with Fungizone (1.25 μg/ml, Gibco, Grand Island, NY). For IFAs, infected cells were fixed in 4% paraformaldehyde 96 hours post infection, washed in 1x PBS and permeabilized for 10 min at RT in 0.1% TritonX-100/PBS and incubated for 15 min in FX Signal Enhancer (Thermofisher #136933). Samples were blocked for 1 hour in 10% BSA/10% goat serum/0.1% TritonX100 and stained mAb 4C9 specific for Pf Hsp70 (REF - F. Zavala, JHU), followed by anti-mouse IgG-Alexa Fluor 488. Images were acquired using a LSM700 laser scanning confocal microscope (Zeiss AxioObserver, Carl Zeiss AG, Oberkochen, Germany) with a 63×/1.4 PlanApo oil objective, and images were acquired using ZEN software (Carl Zeiss AG, Oberkochen, Germany). For live imaging, cells were directly imaged using an inverted Zeiss Axio Observer Z1 microscope with a Yokogawa CSU22 spinning disk using an EMCCD camera and 3i slidebook 5.0 software.

### Gametocyte Assays

Pf-*peg4*-tdTomato gametocytes were induced synchronously according to previous methods (Fivelman et al. Mol Biochem Parasitol 2007) and asexual stages were eliminated by treatment with 50mM GlcNAc (Alfa Aesar, Haverhill, MA) and 20 U/ml heparin (Sigma) for 3 days following re-invasion. On days 2-12 of gametocytogenesis, cultures were stained with 16 μM Hoechst33342 (Invitrogen) and 50 nM DilC1(5) (Invitrogen) for 30 min at 37°C. Mean tdTomato fluorescence was measured using a Cytek DxP12 flow cytometer for >1000 viable gametocytes, gated as infected (Hoechst33342-high), hemozoin-containing (depolarized side scatter-high) RBCs with positive membrane potential (DilC1 (5)-high). Fluorescence images of stained gametocytes were acquired at 1000x magnification using a Leica DM-6000B. tdTomato and Hoechst 33342 Z-stacks were deconvoluted using the non-linear least square algorithm of the DeconvolutionLab2 plug-in of ImageJ and representative slices are shown along with DIC brightfield images. Image of tdTomato-positive microgametes was acquired at 600x magnification after treating cultures of Stage V (day 12) gametocytes with 20 *μ*M Xanthurenic Acid (Sigma) at room temperature.

## Acknowledgements

The authors would like to thank the Bloomberg Family Foundation for their generous support of the Insectary and Parasitology core facilities at the Johns Hopkins Malaria Institute. We also thank Chris Kizito for expert mosquito rearing and the Johns Hopkins School of Medicine Microscopy Facility (MicFac). This work was supported by the National Institutes of Health (R01 AI132359 to PS and F31 pre-doctoral fellowship F31AI136405 to CTH), the Bill & Melinda Gates Foundation (OPP1040399 to DAF), the Johns Hopkins Malaria Research Institute (post-doctoral fellowship to CSH and pre-doctoral fellowship to PCD), the Swedish Research Council (JA) and the Swedish Society of Medicine (JA).

---

**Supplemental Figure.**
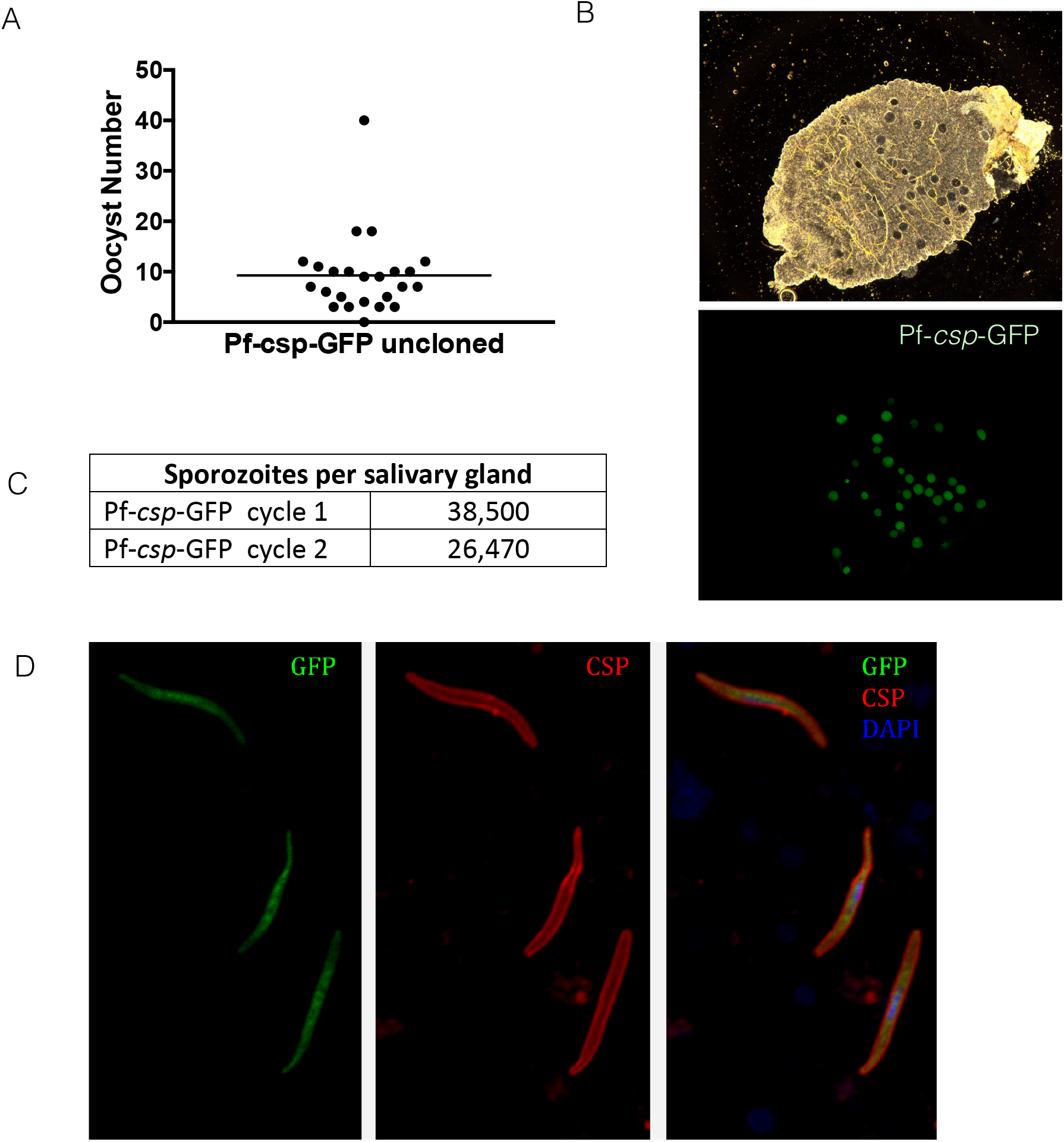
Passage through the mosquito of uncloned Pf-*csp*-GFP line is normal. A) Midgut oocyst numbers and salivary gland sporozoite numbers from two mosquito cycles are shown. Oocysts were counted on day 10 post-infectious blood meal and shown are results from 2 independent experiments in which midguts from 24 mosquitoes were dissected and counted. B) Phase and fluorescence image of an infected midgut. Though the line is not cloned, the majority of oocysts are fluorescent. C) Salivary gland sporozoites were counted on day 14 post-infectious blood meal and shown is the mean of salivary gland loads from 20 mosquitoes for each cycle. D) Immunofluorescence image of salivary gland sporozoites, fixed and stained for PfCSP (red) to visualize the plasma membrane of the sporozoite. The endogenous GFP fluorescence appears to be primarily cytoplasmic (green). DNA was stained with DAPI (blue).

